# The Evolution of Larger Size in High Altitude *Drosophila melanogaster* has a Polymorphic Genetic Architecture

**DOI:** 10.1101/2021.10.05.463286

**Authors:** Quentin D. Sprengelmeyer, Justin B. Lack, Dylan T. Braun, Matthew J. Monette, John E. Pool

## Abstract

Important uncertainties persist regarding the genetic architecture of adaptive trait evolution in natural populations, including the number of genetic variants involved, whether they are drawn from standing genetic variation, and whether directional selection drives them to complete fixation. Here, we take advantage of a unique natural population of *Drosophila melanogaster* from the Ethiopian highlands, which has evolved larger body size than any other known population of this species. We apply a bulk segregant quantitative trait locus (QTL) mapping approach to four unique crosses between highland Ethiopian and lowland Zambian populations for both thorax length and wing length. Results indicated a persistently variable genetic basis for these evolved traits (with largely distinct sets of QTLs for each cross), and at least a moderately polygenic architecture with relatively strong effects present. We complemented these mapping experiments with population genetic analyses of QTL regions and gene ontology enrichment analysis, generating strong hypotheses for specific genes and functional processes that may have contributed to these adaptive trait changes.

Finally, we find that the genetic architectures our QTL mapping results for size traits mirror those from similar experiments on other recently-evolved traits in this species. Collectively, these studies suggest a recurring pattern of polygenic adaptation in this species, in which causative variants do not approach fixation and moderately strong effect loci are present.

## Introduction

Well into the genomic era, considerable debate persists over the types of genetic architectures that underlie adaptive evolution. For example, it is unclear how polygenic adaptive phenotypic changes tend to be – genes of major effect on adaptive traits are often reported (*e.g.* Miller et al. 2014; van’t Hof et al. 2016), and in the case of local adaptation, these are more likely to overcome the homogenizing force of migration (Yeaman & Whitlock 2011). However, it is possible that most adaptive events may instead involve large numbers of small-effect changes (Pritchard & Di Rienzo 2010; Rockman 2011). It is also unclear how often adaptive variants are selected as newly occurring mutations (*e.g.* Linnen *et al.* 2009), versus selection on standing genetic variation after an environmental change (*e.g.* Colosimo *et al.* 2005). In the latter case, the detection of “soft sweeps” is a distinct and more challenging exercise than for classic “hard sweeps” (Pennings & Hermisson 2006). It is also unclear how often adaptive variants actually reach fixation, versus remaining polymorphic due to factors such as traits reaching a new optimum or threshold value, changes in selective pressures, balanced equilibria such as heterozygote advantage, or ongoing migration (Stephan 2016; Höllinger et al. 2019; Thornton 2019; Barghi & Schlötterer 2020; Barghi et al. 2020; Stephan & John 2020).

Population genomic scans for natural selection provide some insight into the genetic basis of adaptive evolution, identifying large numbers of loci with signals of recent positive selection, and estimating the frequency at which different functional categories of sites are targeted. However, the biological basis of natural selection at these loci is usually not clear from genetic variation alone, and the properties of adaptive mutations may depend on the biological process (*e.g.* morphological vs. physiological changes; Carroll 2008; Liao et al. 2010). Therefore, an essential complement to population genomic scans is detailed experimental case studies of the genetic basis of specific adaptive phenotypic changes, in order to gain a clearer and more nuanced understanding of how natural selection operates at the genetic level.

The molecular and evolutionary genetics model *Drosophila melanogaster* provides an efficient system for illuminating the genetic basis of evolutionary change, in part because of its ease of laboratory study, its well-developed molecular genetic toolkit, and its compact and well-annotated genome. *D. melanogaster* expanded from a warm ancestral range in southern-central Africa to occupy diverse worldwide environments (Sprengelmeyer et al. 2020). Latitude and especially altitude gradients allow the comparison of geographically proximate, closely related populations from contrasting environments. Phenotypic differences between genetically similar populations provide ideal raw material for studies of evolution at the genetic level, because the power of population genetic scans for local selection is maximized, and once the relevant genes are identified, the number of plausible causative mutations that differ between populations may be limited.

Size is a fundamental organismal quality. In *D. melanogaster* and other Drosophilids, larger body size is correlated with cooler latitudes (David et al. 1977; Gilchrist & Partridge 1999) and may provide a fitness advantage in cool environments (McCabe & Partridge 1997; Reeve et al. 2000; Bochdanovits & De Jong 2003). Instead of a direct effect of size on thermal tolerance (*Drosophila* are small enough to be virtually isothermic with their environment), higher larval density in the tropics may select for earlier pupation, leading to smaller adults, while in cooler regions viability selection may favor larger, more robust adults (Partridge & French 1996).

In other Drosophilid species, larger flies are also found at higher altitudes (Stalker & Carson 1948; Norry et al. 2001), but this phenomonen was little studied in *D. melanogaster* until recently (Louis et al. 1982). In the past decade, a unique highland Ethiopian population of *D. melanogaster* was found to be the largest known naturally-occurring members of this species, with particularly enlarged wings (Pitchers et al. 2013; Klepsatel et al. 2013; Klepsatel et al. 2014; Fabian et al. 2015; Lack et al. 2016a; Lack et al. 2016b). The increase in wing size is associated with lower wing loading, which can improve flight performance (*e.g.* Petavy et al. 1997) and may benefit flies in highland environments that are persistently cool (limiting the speed of wing movement) and feature thinner air (providing less resistance against fly wings).

Comparing wing length between a highland Ethiopian population and a low altitude ancestral range population from Zabmia, phenotypic differentiation (*Q_ST_* = 0.985) greatly exceeded genetic differentiation (genome-wide *F_ST_* = 0.151), implying that directional selection acted on wing length or a pleiotropically correlated trait (Lack et al. 2016a). The species is only estimated to have occupied the Ethiopian highlands about 2,700 years ago (Sprengelmeyer et al. 2020), or roughly 40,000 fly generations ago (based on 15 generations per year; Turelli & Hoffmann 1995; Pool 2015). In light of an effective population size on the order of one million for this lineage (Sprengelmeyer et al. 2020), the evolution of larger size has occurred on a recent population genetic time scale (~0.01 autosomal coalescent units).

There has been some progress on understanding the tradeoffs and mechanisms involved in this population’s size evolution. Compared to a low altitude Zambian population from the ancestral range, Ethiopian flies lay fewer but larger eggs, which develop into larger adults without prolonging the larval growth phase (Lack et al. 2016b). Ethiopian size changes were found to involve increases in cell size (likely a function of increased somatic ploidy; Smith and Orr-Weaver 1991; Edgar and Orr-Weaver 2001) as well as cell proliferation (Lack et al. 2016b). The evolution of larger wings in Ethiopian *D. melanogaster* was accompanied by a decanalization of wing development, implying that ancestral buffering mechanisms had been disrupted in the course of adaptive trait evolution (Lack et al. 2016a).

The genetic basis of Ethiopian size evolution has not been investigated. Outside Africa, initial progress has been made to identify genes underlying latitude-size clines in *D. melanogaster* outside Africa. In Australia, *Dca* and *srp* are potential contributors to wing and body size differences, respectively (Lee et al. 2011; Chen et al. 2012). Association testing (Jumbo-Lucioni et al. 2010) and experimental evolution (Turner et al. 2011) have suggested that many genes could influence within-population body size variation. However, quantitative trait locus (QTL) mapping of size differences between high and low latitude populations has suggested a few major loci, with uneven chromosomal contributions not predicted by a highly polygenic model (Calboli et al. 2003). Hence, the polygenicity of body size variation may depend on whether diversity is examined within populations where stabilizing selection may predominate, or between populations where adaptive phenotypic evolution is suspected.

In this study we aim to understand the genetic architecture of adaptive trait evolution, using the Ethiopian population’s thorax and wing size changes as model traits. Here, thorax legnth represents a proxy for overall body size, whereas wing length represents a trait that has particularly evolved in this population. We focus on the polygenicity of trait evolution and genetic predictability within a population. We perform bulk segregant analysis to ascertain QTLs that are involved in thorax and wing size trait evolution. We also use population genetic statistics and Gene Ontology enrichment to find evidence of local adaptation and to identify candidate genes for future functional investigation.

## Material and Methods

### Experimental Populations

All flies used in the experiment had been inbred for 8 generations from wild-caught isofemale lines (Lack et al. 2015). The populations came from Fiche, Ethiopia (EF, 9.81° N, 38.63° E, alt. 3070 m) and Siavonga, Zambia (ZI, 16.54° S, 28.72° E, alt. 530 m). These strains were free of any common polymorphic inversions. All flies used were raised at 20° C on medium prepared in batches of 4.5 L water, 500 mL cornmeal, 500 mL molasses, 200 mL yeast, 54 g agar, 20 mL propionic acid, and 45 mL tegosept 10% (in 95% ethanol).

### Bulk Segregant Analysis

To determine what region of the genome harbor the causative variants responsible for the evolution of larger thorax and wing size, bulk segregant analysis was performed to detect quantitative trait loci (QTL). Four different population cages were started with unique strains of smaller thorax and wing size (Zambia) and thorax and wing population (Ethiopia) lines. Each population cage is 28 × 14 × 15 cm and has 14 vials containing the above medium. In each population cage, reciprocal crosses were established between eight inbred parental individuals of each strain (Zambia and Ethiopia). From each reciprocal cross, 125 F1 offspring of each sex were used to establish the second generation. For the duration of the experiment, non-overlapping generations were maintained at ~1200 individuals (Figure 1). Adult flies were allowed to lay eggs on the food for one week before being removed. The food vials were replaced when adult flies in the cage were 7-10 days old. At the 16^th^ generation, 600 3-5 day old female flies from each population cage were measured as described below. For each trait, thorax size and wing size, the flies were placed into pools constituting the 10% smallest (*N*=60) and 10% largest (*N*=60) individuals, with the remaining individuals discarded.

**Figure 1.**
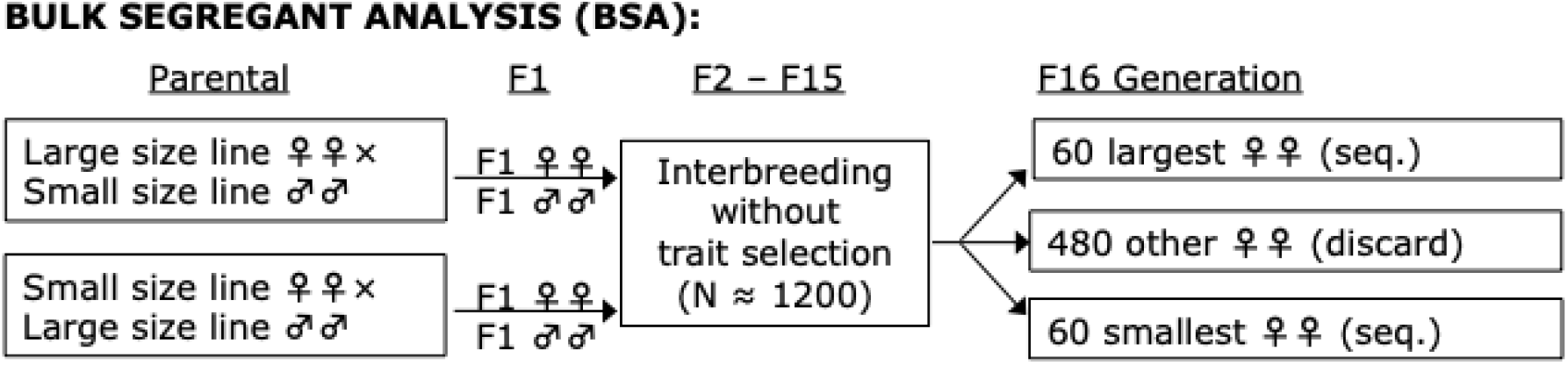
The bulk QTL mapping experimental design is illustrated. As further described in the Materials and Methods, F1 offspring of reciprocal crosses were allowed to interbreed in a relatively large population without selection until the F15 generation, at which point 600 females were sorted to obtain the top and bottom 10% for a size trait for sequencing. This design allows a large number of unique recombination events to take place, which should improve mapping performance.

### Body Size

To measure thorax and wing size, we followed the protocol described in Lack et al. (2016a). Thorax size measurements were in 3-5 day old adult females. From each mapping cross, females were photographed with a digital camera attached to a stereo dissecting microscope (AmScope SM-4BX), and thorax length was measured from the base of the anterior humeral bristle to the posterior tip of the scutellum (Lack et al. 2016a). For wing size, we also examined 3-5 day old adult females from each of the mapping crosses. For five females per cross, a wing was removed and photographed at 509 magnification using a digital camera attached to a compound microscope (Olympus BH-2). The length and depth of each wing were then measured using ImageJ version 1.48 (http://imagej.nih.gov/ij/), we measured a straight line drawn from the intersection of the anterior crossvein and L4 longitudinal vein, to where the L3 longitudinal vein intersects the wing margin. For depth, we measured a straight line from the intersection of the L5 longitudinal vein and the posterior wing margin, passing through the intersection of the posterior crossvein and L4, and terminating at the anterior wing margin. For wing area, we imaged individual wings using the “wing grabber” apparatus described by Houle et al. (2003), and wing area was determined by outlining each wing using ImageJ version 1.48 (http://imagej.nih.gov/ij/), and the reported area for each cross is the mean of the five wings.

### Genome preparation

We sequenced the genomes of pooled samples (N=30 individuals) for the parental lines and two such pools for each of the large- and small-size groups (0-5% and 5-10% extremes for each direction, summing to N=60 total for each extreme). Genomic DNA was obtained using a chloroform extraction and ethanol precipitation protocol. The DNA was fragmented with a Bioruptor sonicator (Diagenode), and paired-end libraries with ~300 bp inserts prepared using NEBNext DNA Library Prep Reagent Set for Illumina (New England Biolabs no. E6000L). Each library’s concentration and quality was analysed with an Agilent 2100 Bioanalyzer (Agilent Technologies, Inc.). The prepared libraries were sequenced at UW-Madison Biotechnology Center on the Illumina HiSeq 2000 platform. Having concluded that the full 10% extremes would best be analyzed together (Pool 2016), we merged reads from the 0-5% and 5-10% pools (similar numbers of reads were obtained from these pools in each case) before proceeding with the analysis.

### Genome alignment

All the raw data that passed the Illumina filters were processed using a Perl-scripted pipeline. Reads from each sequenced genome were mapped to the *D. melanogaster* reference genome (release 5.57) obtained from Flybase (www.flybase.org), with the default parameters in BWA ver. 0.6.2-r126 (Li and Durbin 2009). Using Stampy ver. 1.0.21(Lunter and Goodson 2011), the BAM files were then remapped. With samtools ver. 0.1.18 (Li et al. 2009) reads were filtered for a mapping quality of 20 and for proper pairs. The BAM files were further processed by removing unmapped reads and sorted by coordinate, and PCR duplicates were marked using Picard ver. 1.109 (http://picard.sourceforge.net). To improve the alignment around indels we used GATK ver. 3.2 (McKenna et al. 2010). The average depth of coverage per genome was calculated for the parental lines and the low and high tolerant lines (Table S1).

### Quantitative Trait Locus (QTL) Mapping

Synchronised mpileup files for the aligned genomes were created with the PoPoolation2 ver. 1.201 software package (Kofler et al. 2011). The two large (and two small) pools from a given cross were then combined with a custom perl script. Ancestry difference (*a_d_*) was then calculated with each biallelic SNP (Bastide et al. 2016). Ancestry difference estimates the difference between the proportion of the large-fly pool’s sequencing reads carrying an allele from the large (Ethiopia) parental line and that same proportion from the small-fly pool. It was estimated as:

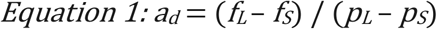

Where *p_L_* is the frequency of the major allele in the large parent, *p_S_* is the small parental allele, *f_L_* is the frequency of the large parent allele in the large pool of F16 offspring, and *f_S_* is that same allele’s frequency in small F16 offspring. The five chromosomal arms (X, 2L, 2R, 3L, and 3R) were divided into windows based on SNP density (Lack et al. 2015) which created 2728, 3131, 2357, 2956, and 2935 windows respectively, each roughly 8.4-kb in size on average. Only sites that had a parental strain frequency difference of ≥ 0.25 were used in the analysis. A simulation-based inference for BSA mapping (SIBSAM) was performed (Pool 2016) to identify significant QTLs and calculate their confidence intervals and effect sizes. The scripts used for SIBSAM can be found at: http://github.com/JohnEPool/SIBSAM1. SIBSAM is able to evaluate both primary QTL peaks and flanking secondary QTL peaks, evaluating whether ragged peaks contain significant evidence for more than one QTL. Forward simulations incorporate recombination in multiple individuals for multiple generations, selection on phenotype in the final generation with additivity, plus environmental variance, and then the sampling of sequence reads to obtain *a_d_*.

### Genetic differentiation and Gene Ontology (GO) enrichment analysis

QTLs identified in the previous step will contain many genes that may or may not be involved in the evolution of these traits. To help identify the causative genes within the significant QTLs for thorax and wing size, window *F_ST_* and maximum SNP *F_ST_* per window (“SNP *F_ST_*”), and the haplotype statistic *χ_MD_* (Lange & Pool 2016) were analyzed. Genomes from Zambia (n=197) and Ethiopia (n=68) were used from the *Drosophila* Genome Nexus (Lack et al. 2015). The *χ_MD_* compares length of identical haplotype blocks among individuals in one population versus another. The comparisons are made within each of the five chromosomal arms (X, 2L, 2R, 3L, and 3R), which were divided into windows based on SNP density (Lack et al. 2015) which created 2728, 3131, 2357, 2956, and 2935 windows respectively each roughly 8.4-kb in size on average. To narrow down potential candidate genes, a chromosomal arm quantile outlier approach was used to identify genes with an extreme population genetic signal. We classified outlier regions windows that were in the top 2.5% quantile in any of the three statistics. In order to form an outlier region, a maximum of two non-outlier windows are allowed between two outlier windows. Genes associated with outlier windows (overlapping them or the nearest gene in either direction) were retained for subsequent analysis.

We preformed a gene ontology (GO) enrichment analysis to identify potential functional categories that may contribute to the contrasting phenotypes found between the Zambia and Ethiopia populations. The outlier genes that were identified in the significant QTL regions were used for window-based GO enrichment analysis (Pool et al. 2012). A GO enrichment analysis was conducted for both thorax and wing size. A *P* value was calculated based on the probability of observing a given number of outlier genes from a GO category. *P* values were obtained from permutation in which outlier regions were randomly reassigned 10,000 times.

## Results

### Quantitative Trait Locus (QTL) Mapping

We used bulk segergant analysis to perform QTL mapping for both thorax and wing length using 4 different unique between-population crosses. Each mapping population used individual inbred strains from an ancestral range Zambia population smaller thorax and wing length, and from the high altitude Ethiopia population that has evolved larger thorax and wing length. In our bulk segergant analysis, offspring of reciprocal crosses were allowed to interbreed for 16 non-overlapping generations without selection at a large population size (N ≈ 1,200). After the 16^th^ generation, 600 adult females were measured for both thorax and wing length and the top and bottom 10% of individuals were grouped for pooled genomic sequencing (Figure 1; Materials and Methods). SIBSAM (Pool 2016) was then used to identify primary and secondary QTL peaks, along with their estimated effect sizes and genomic confidence intervals.

### QTL mapping

For thorax length, four Ethiopia × Zambia mapping crosses revealed a total of 12 significant peaks (Figure 2; Table S2). The EF8N cross had one significant peak with an estimated effect size of ~17%. EF15N had two significant peaks, each having an estimated effect size of ~15%. EF73N had the most significant peaks with a total of five, and these had estimated effect sizes that ranged between 12% and 20%. EF86N had four significant peaks with estimated effect sizes between 13% and 16%.

**Figure 2.**
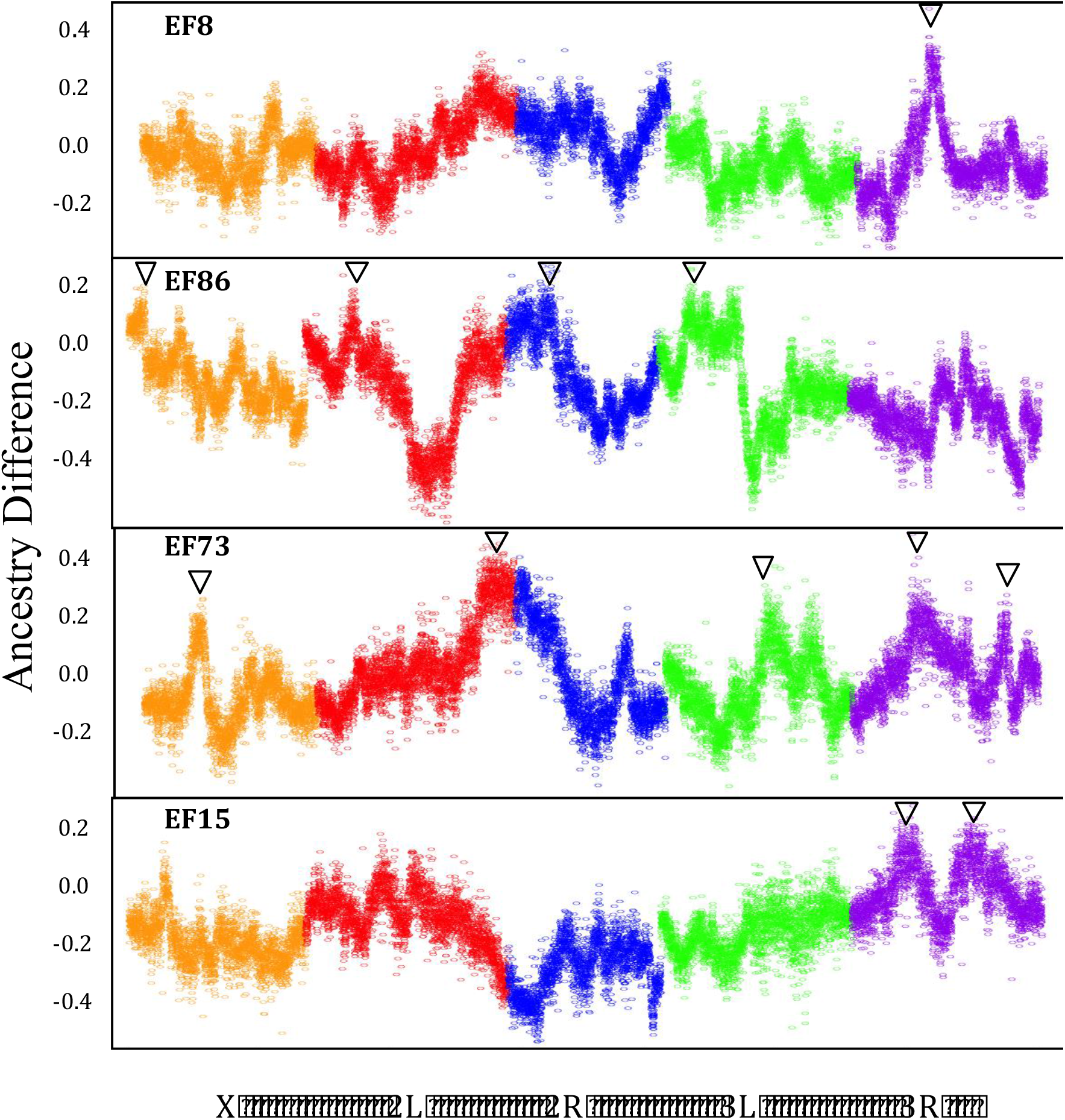
Significant QTL peaks for four Ethiopia/Zambia thorax length crosses. A point for each ~8 kb window corresponds to the average difference in ancestry from the larger parental strain between the large and small F16 pools (y-axis). Significant primary or secondary QTL peaks are denoted with an arrow. The significance threshold for primary peaks is approximately 0.17.

For wing length, these same four crosses revealed a total of 33 significant peaks (Figure 3; Table S3). EF8N had a total of twelve significant peaks, with estimated effect sizes that ranged between 7% and 24%. EF15N had 3 significant peaks, with estimated effect sizes that range between 16% and 24%. EF73N had a total of 10 significant peaks, with estimated effect sizes that ranged between 6% and 27%. EF86N had 8 significant peaks, with estimated effect sizes that ranged between ~11%-25%.

**Figure 3.**
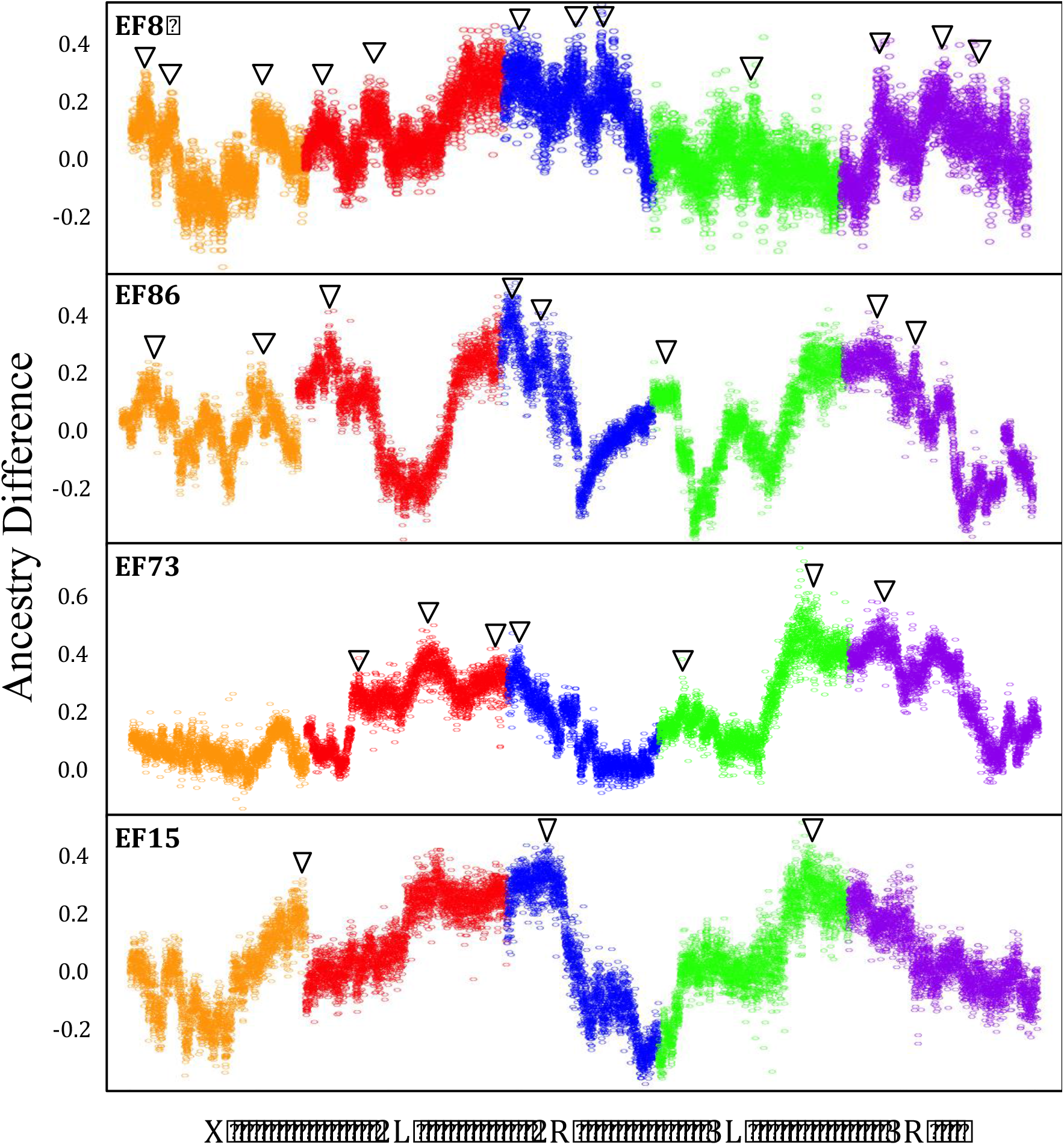
Significant QTL peaks for four Ethiopia/Zambia wing length crosses. A point for each ~8 kb window corresponds to the average difference in ancestry from the larger parental strain between the large and small F16 pools (y-axis). Significant primary or secondary QTL peaks are denoted with an arrow. The significance threshold for primary peaks is approximately 0.17.

In general, very different QTL landscapes were observed between independent Ethiopia/Zambia crosses (Figure 3). In some cases, QTLs do overlap between crosses, which may reflect either chance (different QTLs located close together) or else genuine sharing of causative variants underlying thorax and/or wing size. We identified QTL overlap when the QTL peak of one cross overlaps with the genomic confidence interval of another cross. For thorax length there are no regions between the four Ethiopia crosses where a QTL peak overlapped with another peak’s genomic confidence interval (Figure 4). However, for wing length between the four crosses there were 12 regions where QTL peaks overlapped with genomic confidence intervals involving 12 of the 33 QTLs (Figure 4). Within these overlapping peaks there are no overlap between all four crosses.

**Figure 4.**
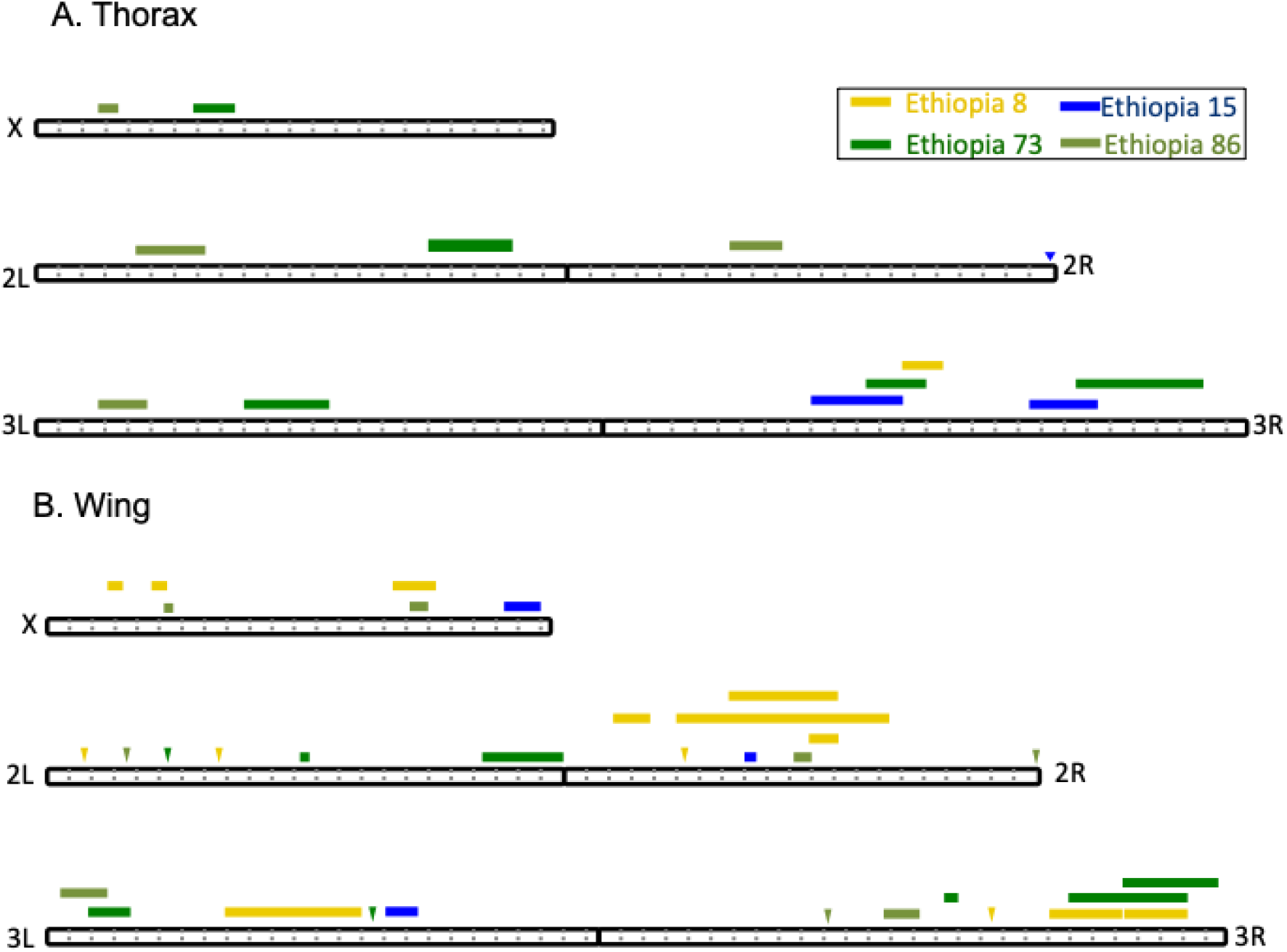
The locations of significant QTLs on the five euchromatic chromosome arms of *D. melanogaster*. The colors indicate for four Ethiopia strains used in mapping crosses for **A**) thorax length and **B**) wing length. The width of each box indicates the 90% C.I. of each QTL. Intervals that are less than 10 kb in width are marked with triangles. Dotted gray lines indicate Mb increments.

Some differences in the significant QTLs between crosses could represent chance detection of a shared QTL in some crosses but not others. However, with this experimental design we expect to have >90% power to detect a QTL with 20% effect size (Pool 2016). Hence, at least for several of the strongest of the QTLs detected here, their absence in other crosses is likely to reflect real differences in genetic architecture.

### Potential Targets of Local Adaptation Within QTL Regions

Regions of the genome where the Zambia and Ethiopia populations greatly differ in their genetic variation may harbor genes involved in these adaptive traits. We used three population genetic statistics, window *F_ST_*, maximum SNP *F_ST_* within a window, and the haplotype statistic *χ_MD_* to identify possible candidate genes for body and wing size evolution within the significant QTLs. Using three different statistics is advantageous due to the differing power each statistic has in detecting local adaptation, depending on whether selective sweeps are complete or incomplete, or hard versus soft (Lange & Pool 2016). A quantile approach was used identify only regions that had one of the three statistics with a quantile below 0.025 (Tables S4 & S5). There are many genes within these outlier regions with no known role in either thorax or wing size. However, there are also genes known to be involved in size regulation.

For thorax length, genes corresponding to QTLs and population genetic outliers that are known to be involved in growth included *ct* (Thumm & Kadowaki 2001), *spi* (Nagaraj et al. 1999), *bbc* (Liu et al. 2014), *msn* (Kadrmas et al. 2004), *RasGAP1* (Dworkin & Gibson 2006), *scyl* (Reiling and Hafen 2004), and *tara* (Bejarano et al., 2008). Of these, *bbc* and *RasGAP1* provide examples of loci with promisingly narrow *F_ST_* peaks at the SNP level (Figure 5), which may merit targeted investigation by future studies. We noted that *RasGAP1* is also within a wing QTL, and is therefore relevant to the analysis described below as well.

**Figure 5.**
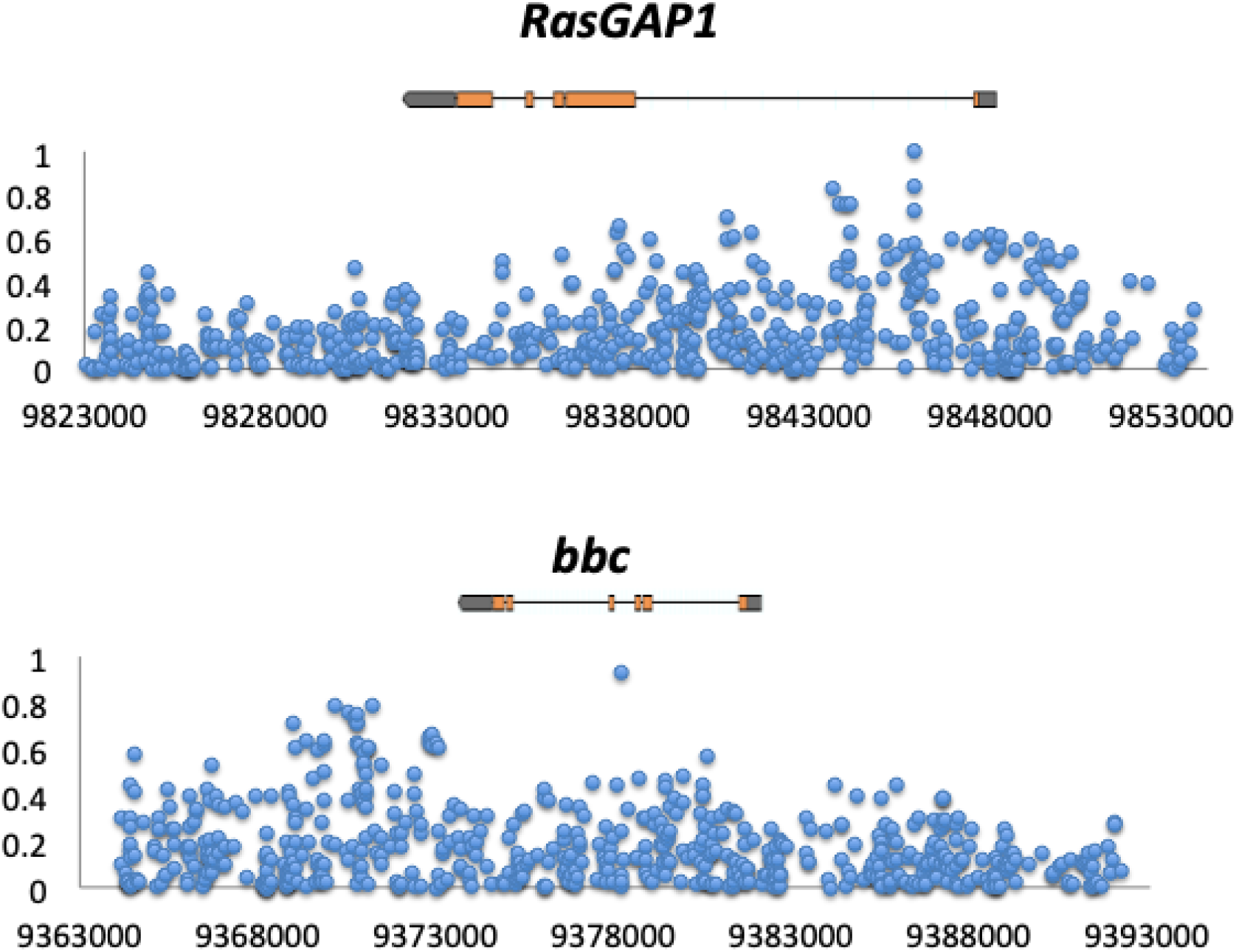
At two candidate genes identified for thorax length evolution (*bbc* and *RasGAP1*), small numbers of SNPs show the highest *F_ST_* values between Ethiopia and Zambia. Depicted above is the gene transcript, while the x-axis indicates bp position along the relevant chromosome arm (release 5).

Within the outlier regions for wing length, these genes included *Dronc* (Verghese et al. 2012*), Dlish* (Wang et al. 2019), *fj* (Villano & Katz 1995), *Pka-C3* (Dworkin & Gibson 2006), *salr* (Wang et al.2017), and *Gbp1* (Koyama and Mirth 2016)*. Dlish* and *Pka-C3* are examples of genes with individual SNPs having high *F_ST_* values (Figure 6). Further functional testing will need to be conducted to establish if genetic variants found within these genes are indeed responsible for the associated phenotypes.

**Figure 6.**
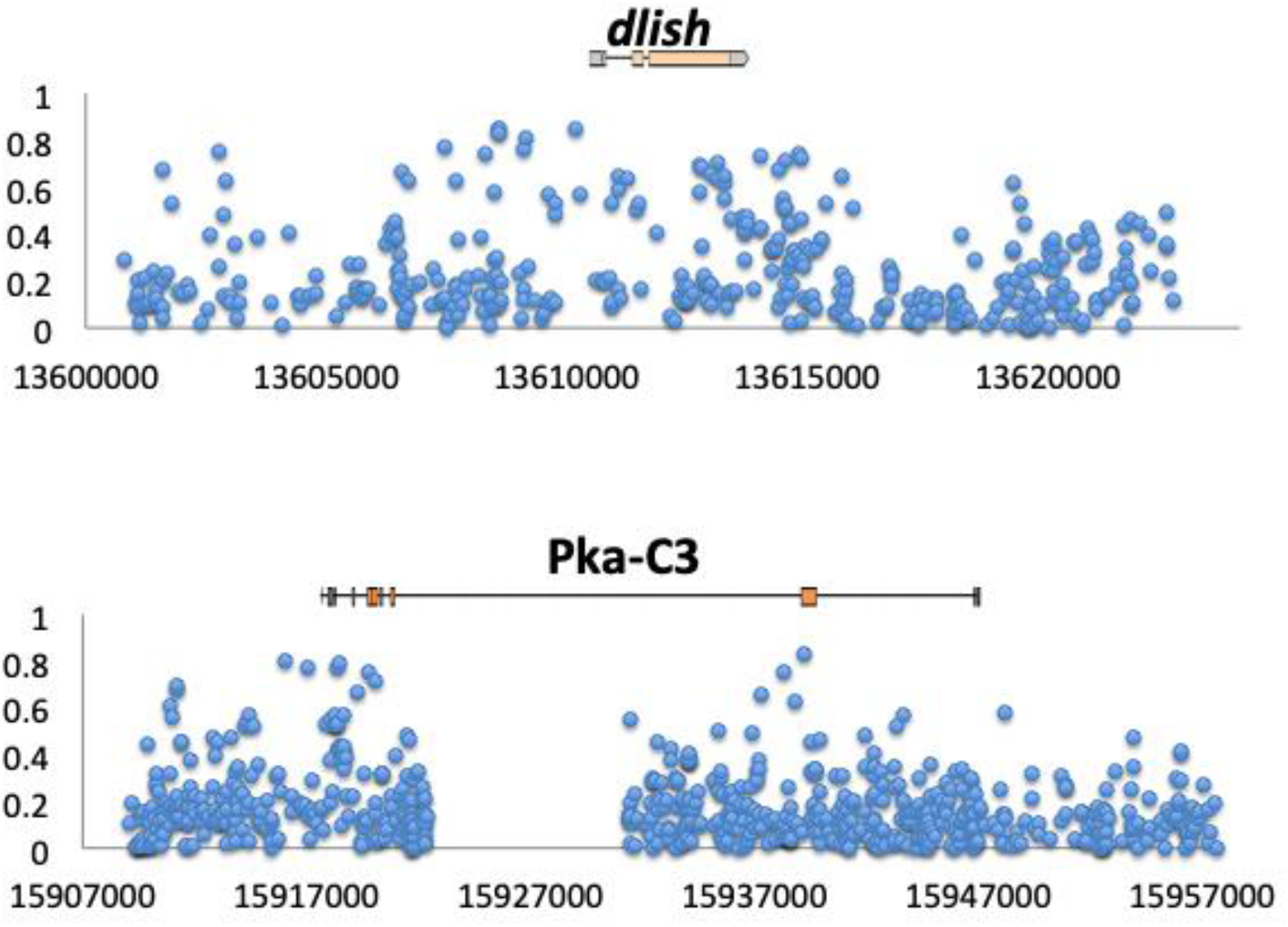
Peaks of SNP *F_ST_* center on two candidate genes identified for wing size evolution (*dlish* and *Pka-C3*), showing elevated genetic differentiation between Ethiopia and Zambia at these genes. Depicted above is the gene transcript, while the x-axis indicates bp position along the relevant chromosome arm (release 5).

### Gene Ontology (GO) Enrichment

We conducted individual GO enrichment analysis for thorax and wing length. We used only the genes found in the outlier windows located within significant QTL regions from the four crosses. Functional categories that yielded raw *P* values below 0.001 are listed in Tables S6 & S7. These included categories either known or potentially involved in body and wing size. For thorax size, the top categories included: negative regulation of Ras protein signal transduction (Prober & Edgar 2002), regulation of protein polymerization (Fernández et al. 2011), and brahma complex (Krupp et al. 2005). For wing size, the top categories included: neurogenesis (Rutledge et al. 1992), ubiquitin protein ligase binding (Cornell et al. 1999), cellular amino acid catabolic process (Zinke et al. 1999), cellular response to anoxia (Heinrich et al 2011), transmembrane transport (Bartscherer et al. 2006), and negative regulation of proteolysis (Lee et al. 2001). Some of these functional processes might underlie Ethiopia size adaptation, while others may be driven by unrelated trait evolution in this high altitude population.

## Discussion

We employed quantitive and population genetic strategies to investigate the genetic architecture of adaptive size evolution in our highland Ethiopia population. Our bulk segergant analysis revealed that between the four crosses, thorax size has 12 associated QTLs with moderate to large effect (~13-20%). However, between the four crosses wing size has 33 QTLs with small to large effects QTLs (~6-27%). A greater ability to detect wing length QTLs than thorax length QTLs may reflect the greater magnitude of the population difference in this trait (Lack et al. 2016b).

One striking result was the lack of QTL overlap between crosses for either thorax or wing size. Between the four thorax crosses there is no overlap. This is especially notable given that we have almost have very high power to detect QTLs with effect size of 20% (Pool 2016) and yet the QTL on chromosome arm 2L with ~20% effect size is not present in any other cross. For wing size, there was overlap in only 12 of the 33 QTL regions and no overlap between all four crosses. The QTLs with the three largest effect sizes of over 25% are present in only one cross. The low QTL overlap between crosses could reflect persistent genetic variation at cauasative loci in the Ethiopian and/or Zambian populations. Given that the Ethiopian population appears to have experienced directional selection for larger size, and still maintains similar genetic variance for size traits as Zambia (Lack et al. 2016b), we suggest that some favored size variants have not reached fixation in the Ethiopian population. There are multiple reasons why favored alleles might not fix, including the Ethiopian population reaching its new optimum or threshold trait value (especially if ample standing variation means that not all large alleles needed to fix), heterozygote advantage, or ongoing adaptation. Indeed, simulation and theory have shown that depending on the genetic architecture of an adaptive trait, non-fixed causative variants may be the norm (Stephan 2016; Höllinger et al. 2019; Thornton 2019; Barghi & Schlötterer 2020; Barghi et al. 2020; Stephan & John 2020).

Our conclusions of persistent variability underlying an evolved trait mirror similar results for pigmentation (Bastide et al. 2016) and for ethanol resistance (Sprengelmeyer et al. 2021) in this same population and others, all from mapping experiments with similar design and scale. With five traits now examined (ethanol resistance, abdominal background color, abdominal stripe width, thorax length, and wing length), consistent patterns are starting to emerge. First, these traits each average at least a few detectable QTLs per cross, with means ranging from 3 to 8.25 (Table 1). Second, there is notably little QTL peak overlap between parallel mapping crosses involving different strains from the same populations. Wing length and ethanol resistance have the most overlap between crosses with ~35% (Table. 1). However, thorax size crosses do not have any overlap. For each of these traits, there are moderately large effect QTLs not present in other crosses. Hence, at the population level, it is fair to say that each of these traits is at least moderately polygenic, and involves non-fixed differences between populations. Third, moderately strong QTLs are consistently present in any given cross, with the average QTL effect size ranging from 13-19% (Table 1), although undetectable smaller effects may be present as well.

**Table 1.** The results of bulk QTL mapping experiments for five different traits. All mapping used the same experimental design described in the Materials and Methods, aside from minor variation in the number of generations of interbreeding (15-20). Both the pigmentation stripe and pigmentation background data is from Bastide et al. (2016), while the ethanol results are from Sprengelmeyer & Pool (2021). Listed are the number of significant QTLs for each mapping population, the proportion of QTLs that overlap between parallel crosses from the same two populations, and the average QTL effect size across all mapping crosses.

| Trait | Avg. QTLs per<br>Cross | Pairwise QTL<br>Overlap | Avg. Effect Size |
| --- | --- | --- | --- |
| Ethanol Resistance | 8 | 35% | 14% |
| Thorax Length | 3 | 0% | 15% |
| Wing Length | 8.25 | 36% | 17% |
| Stripe Width | 3 | 8% | 19% |
| Background Color | 5.67 | 18% | 19% |

Polygenic adaptation may have diverse outcomes, depending in part on the number of segregating variants at the onset of selection that affect a trait, as well as the magnitudes of their effect on the trait relative to the shift in trait optimum. While each of the traits summarized above might be described as “polygenic”, it is worth considering the type of polygenic adaptation that these mapping studies imply. The persistently variable genetic basis of these evolved traits may suggest a scenario of abundant standing genetic variation prior to selection for each of these traits. In light of the consistent presence of moderately strong QTLs for these traits, such standing variation may have included relatively large effect loci, which would experience relatively stronger directional selection during the trait’s evolution. An abundance of standing variation is consistent with the large population size and high genetic diversity of this species (*e.g.* Sprengelmeyer et al. 2020). Further studies will be needed to quantify the models of polygenic adaptation that experiments such as ours indicate, and to assess whether such persistent variability is a widespread outcome of trait evolution not only in this species but across the tree of life.

## Acknowledgments

We would like to thank Jeremy Lange and Tiago Ribeiro for their help with bioinformatics and data collection in this project. We also thank the UW-Madison Center for High Throughput Computing (CHTC) for computational resources and assistance. This research was was supported by the National Institutes of Health grants R01 GM111797, R01 GM127480, R35 GM136306, F32 GM106594, and T32 GM007133.

## Data Availability

All raw sequence data has been deposited in the NIH Short Read Archive, with accession numbers given in Table S1.

---

**Table S3.**
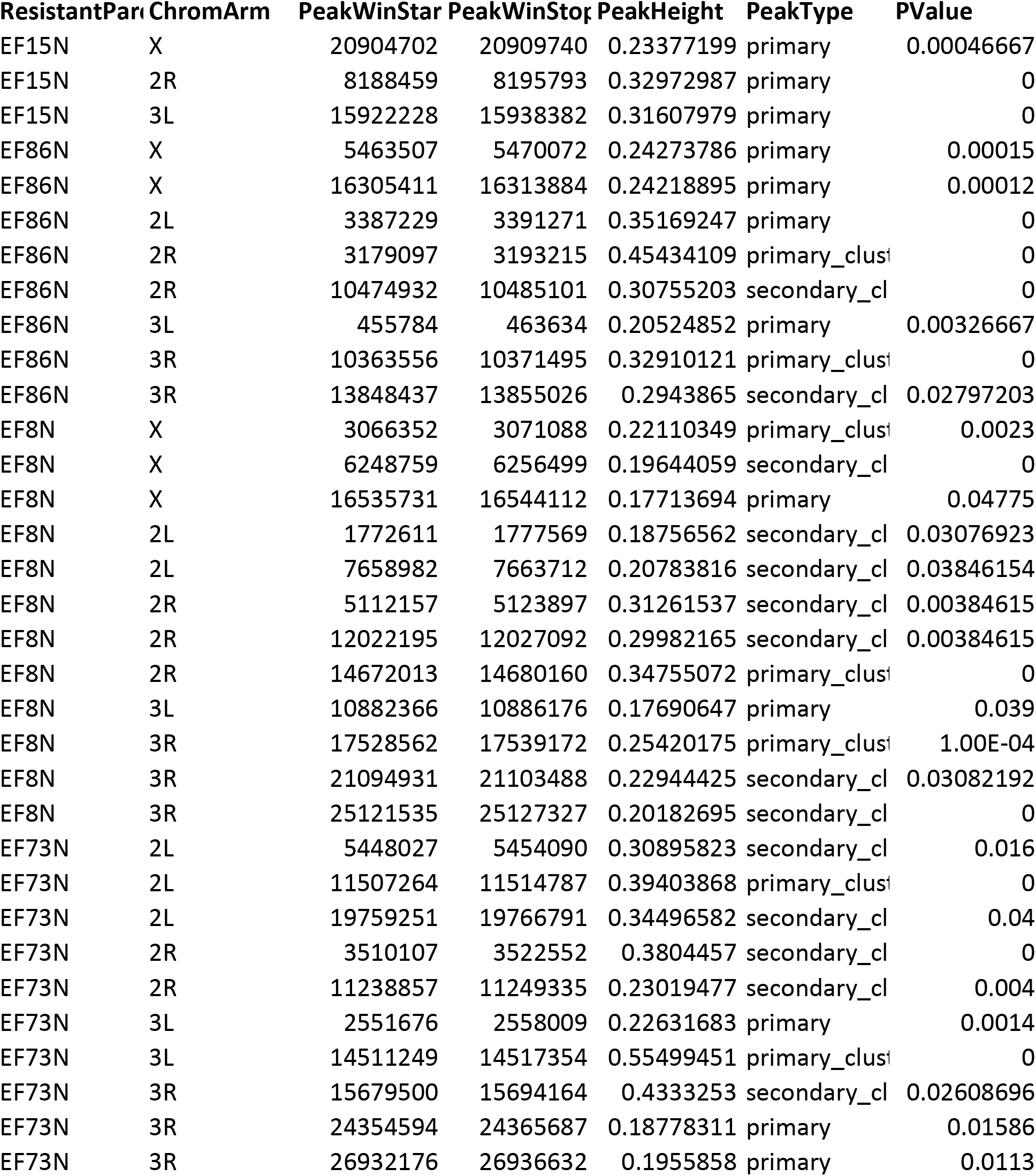
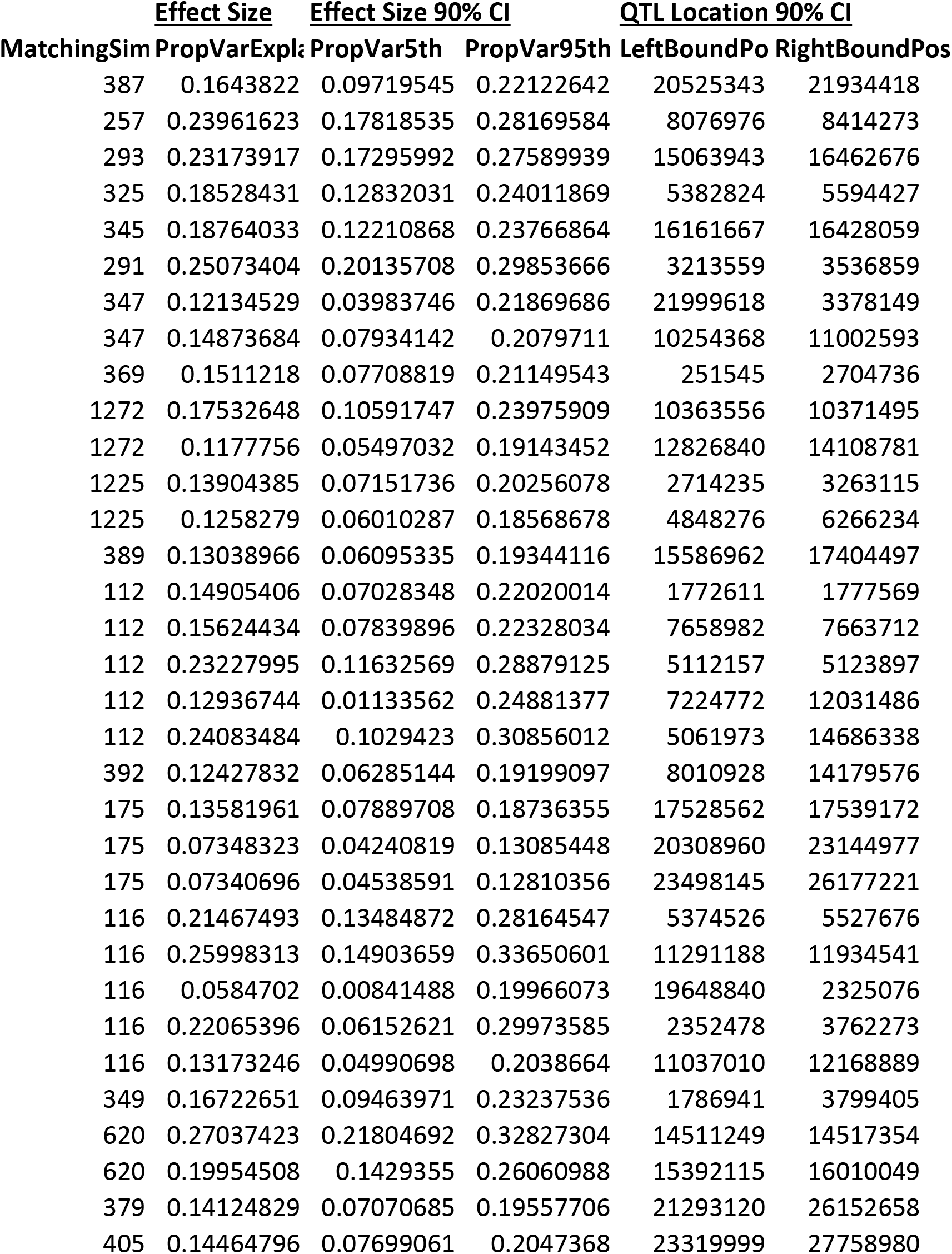
Locations and properties of significant QTLs identified in wing size mapping by SIBSA.

## Literature Cited

Barghi, N., Hermisson, J., & Schlötterer, C. (2020). Polygenic adaptation: A unifying framework to understand positive selection. Nature Reviews Genetics, 21(12), 769–781.

Bartscherer, K., Pelte, N., Ingelfinger, D., & Boutros, M. (2006). Secretion of Wnt Ligands Requires Evi, a Conserved Transmembrane Protein. Cell, 125(3), 523–533.

Bastide, H., Yassin, A., Johanning, E. J., & Pool, J. E. (2014). Pigmentation in *Drosophila melanogaster* reaches its maximum in Ethiopia and correlates most strongly with ultra-violet radiation in sub-Saharan Africa. BMC Evolutionary Biology, 14, 179.

Bastide, H., Lange, J. D., Lack, J. B., Yassin, A., & Pool, J. E. (2016). A variable genetic architecture of melanic evolution in *Drosophila melanogaster*. Genetics, 204(3), 1307–1319.

Bejarano, F., Luque, C. M., Herranz, H., Sorrosal, G., Rafel, N., Pham, T. T., & Milán, M. (2008). A Gain-of-Function Suppressor Screen for Genes Involved in Dorsal–Ventral Boundary Formation in the *Drosophila* Wing. Genetics, 178(1), 307–323.

Bochdanovits, Z., & De Jong, G. (2003). Temperature dependence of fitness components in geographical populations of *Drosophila melanogaster*: changing the association between size and fitness. Biological Journal of the Linnean Society, 80(4), 717–725

Calboli, F. C. F., Kennington, W. J., & Partridge, L. (2003). Qtl Mapping Reveals a Striking Coincidence in the Positions of Genomic Regions Associated with Adaptive Variation in Body Size in Parallel Clines of *Drosophila melanogaster* on Different Continents. Evolution, 57(11), 2653–2658.

Carroll, S. B. (2008). Evo-Devo and an Expanding Evolutionary Synthesis: A Genetic Theory of Morphological Evolution. Cell, 134(1), 25–36.

Chen, Y., Lee, S. F., Blanc, E., Reuter, C., Wertheim, B., Martinez-Diaz, P., Hoffmann, A. A., & Partridge, L. (2012). Genome-Wide Transcription Analysis of Clinal Genetic Variation in *Drosophila*. PLOS ONE, 7(4), e34620.

Colosimo, P. F., Hosemann, K. E., Balabhadra, S., Villarreal, G., Dickson, M., Grimwood, J., Schmutz, J., Myers, R. M., Schluter, D., & Kingsley, D. M. (2005). Widespread Parallel Evolution in Sticklebacks by Repeated Fixation of Ectodysplasin Alleles. Science, 307(5717), 1928–1933.

Cornell, M., Evans, D. a. P., Mann, R., Fostier, M., Flasza, M., Monthatong, M., Artavanis-Tsakonas, S., & Baron, M. (1999). The Drosophila melanogaster Suppressor of deltex Gene, a Regulator of the Notch Receptor Signaling Pathway, Is an E3 Class Ubiquitin Ligase. Genetics, 152(2), 567–576.

David, J. R., Bocquet, C., & De Scheemaeker-Louis, M. (1977). Genetic latitudinal adaptation of *Drosophila melanogaster*: new discriminative biometrical traits between European and equatorial African populations. Genet. Res., 30, 247–255.

de Jong, G., & Bochdanovits, Z. (2003). Latitudinal clines inDrosophila melanogaster: Body size, allozyme frequencies, inversion frequencies, and the insulin-signalling pathway. Journal of Genetics, 82(3), 207–223.

Dworkin, I., & Gibson, G. (2006). Epidermal Growth Factor Receptor and Transforming Growth Factor-β Signaling Contributes to Variation for Wing Shape in *Drosophila melanogaster*. Genetics, 173(3), 1417–1431.

Edgar, B. A., & Orr-Weaver, T. L. (2001). Endoreplication Cell Cycles: More for Less. Cell, 105(3), 297–306.

Fabian, D. K., Lack, J. B., Mathur, V., Schlötterer, C., Schmidt, P. S., Pool, J. E., & Flatt, T. (2015). Spatially varying selection shapes life history clines among populations of *Drosophila melanogaster* from sub-Saharan Africa. Journal of Evolutionary Biology, 28(4), 826–840.

Fernández, B. G., Gaspar, P., Brás-Pereira, C., Jezowska, B., Rebelo, S. R., & Janody, F. (2011). Actin-Capping Protein and the Hippo pathway regulate F-actin and tissue growth in *Drosophila*. Development, 138(11), 2337–2346.

Gilchrist, A. Stuart, & Partridge, L. (1999). A Comparison of the Genetic Basis of Wing Size Divergence in Three Parallel Body Size Clines of *Drosophila melanogaster*. Genetics, 153(4), 1775–1787.

Heinrich, E. C., Farzin, M., Klok, C. J., & Harrison, J. F. (2011). The effect of developmental stage on the sensitivity of cell and body size to hypoxia in *Drosophila melanogaster*. Journal of Experimental Biology, 214(9), 1419–1427.

Hermisson, J., & Pennings, P. S. (2005). Soft Sweeps: Molecular Population Genetics of Adaptation From Standing Genetic Variation. Genetics, 169(4), 2335–2352.

James, A. C., Azevedo, R. B., & Partridge, L. (1995). Cellular basis and developmental timing in a size cline of *Drosophila melanogaster*. Genetics, 140(2), 659–666.

Höllinger, I., Pennings, P. S., & Hermisson, J. (2019). Polygenic adaptation: From sweeps to subtle frequency shifts. PLOS Genetics, 15(3), e1008035.

John, S., & Stephan, W. (2020). Important role of genetic drift in rapid polygenic adaptation. Ecology and Evolution, 10(3), 1278–1287.

Jumbo-Lucioni, P., Ayroles, J. F., Chambers, M. M., Jordan, K. W., Leips, J., Mackay, T. F., & De Luca, M. (2010). Systems genetics analysis of body weight and energy metabolism traits in *Drosophila melanogaster*. BMC Genomics, 11(1), 297.

Kadrmas, J. L., Smith, M. A., Clark, K. A., Pronovost, S. M., Muster, N., Yates, J. R., III, & Beckerle, M. C. (2004). The integrin effector PINCH regulates JNK activity and epithelial migration in concert with Ras suppressor 1. Journal of Cell Biology, 167(6), 1019–1024.

Klepsatel, P., Gáliková, M., Maio, N. D., Huber, C. D., Schlötterer, C., & Flatt, T. (2013). Variation in Thermal Performance and Reaction Norms Among Populations of *Drosophila melanogaster*. Evolution, 67(12), 3573–3587.

Klepsatel, P., Gáliková, M., Huber, C. D., & Flatt, T. (2014). Similarities and Differences in Altitudinal Versus Latitudinal Variation for Morphological Traits in *Drosophila melanogaster*. Evolution, 68(5), 1385–1398.

Kofler, R., Pandey, R. V., & Schlötterer, C. (2011). PoPoolation2: identifying differentiation between populations using sequencing of pooled DNA samples (Pool-Seq). Bioinformatics, 27(24), 3435–3436.

Koyama, T., & Mirth, C. K. (2016). Growth-Blocking Peptides As Nutrition-Sensitive Signals for Insulin Secretion and Body Size Regulation. PLOS Biology, 14(2), e1002392.

Krupp, J. J., Yaich, L. E., Wessells, R. J., & Bodmer, R. (2005). Identification of Genetic Loci That Interact With cut During Drosophila Wing-Margin Development. Genetics, 170(4), 1775–1795.

Lack, J. B., Cardeno, C. M., Crepeau, M. W., Taylor, W., Corbett-Detig, R. B., Stevens, K. A., Langley, C. H., & Pool, J. E. (2015). The Drosophila genome nexus: a population genomic resource of 623 *Drosophila melanogaster* genomes, including 197 from a single ancestral range population. Genetics, 199(4), 1229–1241.

Lack, J. B., Monette, M. J., Johanning, E. J., Sprengelmeyer, Q. D., & Pool, J. E. (2016a). Decanalization of wing development accompanied the evolution of large wings in high-altitude *Drosophila*. Proceedings of the National Academy of Sciences, 113(4), 1014–1019

Lack, J. B., Yassin, A., Sprengelmeyer, Q. D., Johanning, E. J., David, J. R., & Pool, J. E. (2016b). Life history evolution and cellular mechanisms associated with increased size in high-altitude *Drosophila*. Ecology and evolution, 6(16), 5893–5906.

Lack, J. B., Lange, J. D., Tang, A. D., Corbett-Detig, R. B., & Pool, J. E. (2016c). A Thousand Fly Genomes: An Expanded *Drosophila* Genome Nexus. Molecular Biology and Evolution, 33(12), 3308–3313.

Lange, J. D., & Pool, J. E. (2016). A haplotype method detects diverse scenarios of local adaptation from genomic sequence variation. Molecular ecology, 25(13), 3081–3100.

Lee, J. R., Urban, S., Garvey, C. F., & Freeman, M. (2001). Regulated Intracellular Ligand Transport and Proteolysis Control EGF Signal Activation in *Drosophila*. Cell, 107(2), 161–171.

Lee, S. F., Chen, Y., Varan, A. K., Wee, C. W., Rako, L., Axford, J. K., Good, R. T., Blacket, M. J., Reuter, C., Partridge, L., & Hoffmann, A. A. (2011). Molecular Basis of Adaptive Shift in Body Size in *Drosophila melanogaster*: Functional and Sequence Analyses of the Dca Gene. Molecular Biology and Evolution, 28(8), 2393–2402.

Li, H., & Durbin, R. (2009). Fast and accurate short read alignment with Burrows–Wheeler transform. Bioinformatics, 25(14), 1754–1760.

Liao, B.-Y., Weng, M.-P., & Zhang, J. (2010). Contrasting genetic paths to morphological and physiological evolution. Proceedings of the National Academy of Sciences, 107(16), 7353–7358.

Linnen, C. R., Kingsley, E. P., Jensen, J. D., & Hoekstra, H. E. (2009). On the Origin and Spread of an Adaptive Allele in Deer Mice. Science, 325(5944), 1095–1098.

Liu, Y., Wang, W., Shui, G., & Huang, X. (2014). CDP-Diacylglycerol Synthetase Coordinates Cell Growth and Fat Storage through Phosphatidylinositol Metabolism and the Insulin Pathway. PLOS Genetics, 10(3), e1004172.

Louis, J., David, J., Rouault, J., & Capy, P. (1982). Altitudinal variations of Afro-tropical D. melanogaster populations. Dros. Inf. Serv, 58, 100–101.

Lunter, G., & Goodson, M. (2011). Stampy: A statistical algorithm for sensitive and fast mapping of Illumina sequence reads. Genome Research, 21(6), 936–939.

McCabe, J., & Partridge, L. (1997). An Interaction Between Environmental Temperature and Genetic Variation for Body Size for the Fitness of Adult Female *Drosophila* Melanogaster. Evolution, 51(4), 1164–1174.

McKenna, A., Hanna, M., Banks, E., Sivachenko, A., Cibulskis, K., Kernytsky, A., Garimella, K., Altshuler, D., Gabriel, S., Daly, M., & DePristo, M. A. (2010). The Genome Analysis Toolkit: A MapReduce framework for analyzing next-generation DNA sequencing data. Genome Research, 20(9), 1297–1303.

Miller, C. T., Glazer, A. M., Summers, B. R., Blackman, B. K., Norman, A. R., Shapiro, M. D., Cole, B. L., Peichel, C. L., Schluter, D., & Kingsley, D. M. (2014). Modular Skeletal Evolution in Sticklebacks Is Controlled by Additive and Clustered Quantitative Trait Loci. Genetics, 197(1), 405–420.

Norry, F. M., Bubliy, O. A., & Loeschcke, V. (2001). Developmental Time, Body Size and Wing Loading in Drosophila Buzzatii from Lowland and Highland Populations in Argentina. Hereditas, 135(1), 35–40.

Pennings, P. S., & Hermisson, J. (2006). Soft Sweeps III: The Signature of Positive Selection from Recurrent Mutation. PLOS Genetics, 2(12), e186.

Pitchers, W., Pool, J. E., & Dworkin, I. (2013). Altitudinal Clinal Variation in Wing Size and Shape in African *Drosophila melanogaster*: One Cline or Many? Evolution, 67(2), 438–452.

Pool, J. E., Corbett-Detig, R. B., Sugino, R. P., Stevens, K. A., Cardeno, C. M., Crepeau, M. W., Duchen, P., Emerson, J. J., Saelao, P., Begun, D. J., & Langley, C. H. (2012). Population Genomics of Sub-Saharan *Drosophila melanogaster*: African Diversity and Non-African Admixture. PLOS Genetics, 8(12), e1003080.

Pool, J. E. (2015). The mosaic ancestry of the *Drosophila* genetic reference panel and the D. melanogaster reference genome reveals a network of epistatic fitness interactions. Molecular biology and evolution, 32(12), 3236–3251.

Pool, J. E. (2016). Genetic mapping by bulk segregant analysis in *Drosophila*: experimental design and simulation-based inference. Genetics, 204(3), 1295–1306.

Pool, J. E., Braun, D. T., & Lack, J. B. (2017). Parallel evolution of cold tolerance within *Drosophila melanogaster*. Molecular biology and evolution, 34(2), 349–360.

Pritchard, J. K., & Di Rienzo, A. (2010). Adaptation –not by sweeps alone. Nature Reviews Genetics, 11(10), 665–667.

Prober, D. A., & Edgar, B. A. (2002). Interactions between Ras1, dMyc, and dPI3K signaling in the developing *Drosophila* wing. Genes & Development, 16(17), 2286–2299.

Reeve, M. W., Fowler, K., & Partridge, L. (2000). Increased body size confers greater fitness at lower experimental temperature in male *Drosophila melanogaster*. Journal of Evolutionary Biology, 13(5), 836–844.

Reiling, J. H., & Hafen, E. (2004). The hypoxia-induced paralogs *Scylla* and *Charybdis* inhibit growth by down-regulating S6K activity upstream of TSC in *Drosophila*. Genes & Development, 18(23), 2879–2892.

Rockman, M. V. (2012). The QTN Program and the alleles that matter for evolution: All that’s gold does not glitter. Evolution, 66(1), 1–17.

Rutledge, B. J., Zhang, K., Bier, E., Jan, Y. N., & Perrimon, N. (1992). The *Drosophila spitz* gene encodes a putative EGF-like growth factor involved in dorsal-ventral axis formation and neurogenesis. Genes & Development, 6(8), 1503–1517.

Smith, A. V., & Orr-Weaver, T. L. (1991). The regulation of the cell cycle during Drosophila embryogenesis: The transition to polyteny. Development, 112(4), 997–1008.

Sprengelmeyer, Q. D., Mansourian, S., Lange, J. D., Matute, D. R., Cooper, B. S., Jirle, E. V., Stensmyr, M. C., & Pool, J. E. (2020). Recurrent collection of *Drosophila melanogaster* from wild African environments and genomic insights into species history. Molecular Biology and Evolution 37(3), 627–638.

Sprengelmeyer, Q. D., & Pool, J. E. (2021). Ethanol resistance in *Drosophila melanogaster* has increased in parallel cold-adapted populations and shows a variable genetic architecture within and between populations. Ecology and Evolution, Accepted.

Stalker, H. D., & Carson, H. L. (1948). An Altitudinal Transect of *Drosophila robusta* Sturtevant. Evolution, 2(4), 295–305

Stephan, W. (2016). Signatures of positive selection: From selective sweeps at individual loci to subtle allele frequency changes in polygenic adaptation. Molecular Ecology, 25(1), 79–88.

Thornton, K. R. (2019). Polygenic Adaptation to an environmental shift: Temporal dynamics of variation under gaussian stabilizing selection and additive effects on a single trait. Genetics, 213(4), 1513–1530.

Thumm, M., & Kadowaki, T. (2001). The loss of *Drosophila* APG4/AUT2 function modifies the phenotypes of cut and Notch signaling pathway mutants. Molecular Genetics and Genomics, 266(4), 657–663.

Turner, T. L., Stewart, A. D., Fields, A. T., Rice, W. R., & Tarone, A. M. (2011a). Population-Based Resequencing of Experimentally Evolved Populations Reveals the Genetic Basis of Body Size Variation in *Drosophila melanogaster*. PLOS Genetics, 7(3), e1001336.

van’t Hof, A. E., Edmonds, N., Dalíková, M., Marec, F., & Saccheri, I. J. (2011). Industrial melanism in British peppered moths has a singular and recent mutational origin. Science, 332(6032), 958–960.

Verghese, S., Bedi, S., & Kango-Singh, M. (2012). Hippo signalling controls Dronc activity to regulate organ size in *Drosophila*. Cell Death & Differentiation, 19(10), 1664–1676.

Villano, J. L., & Katz, F. N. (1995). Four-jointed is required for intermediate growth in the proximal-distal axis in *Drosophila*. Development, 121(9), 2767–2777.

Wang, D., Li, J., Liu, S., Zhou, H., Zhang, L., Shi, W., & Shen, J. (2017). Spalt is functionally conserved in *Locusta* and *Drosophila* to promote wing growth. Scientific Reports, 7(1), 44393.

Wang, X., Zhang, Y., & Blair, S. S. (2019). Fat-regulated adaptor protein Dlish binds the growth suppressor Expanded and controls its stability and ubiquitination. Proceedings of the National Academy of Sciences, 116(4), 1319–1324.

Yeaman, S., & Whitlock, M. C. (2011). The genetic architecture of adaptation under migration–selection balance. Evolution: International Journal of Organic Evolution, 65(7), 1897–1911.

Zinke, I., Kirchner, C., Chao, L. C., Tetzlaff, M. T., & Pankratz, M. J. (1999). Suppression of food intake and growth by amino acids in *Drosophila*: The role of pumpless, a fat body expressed gene with homology to vertebrate glycine cleavage system. Development, 126(23), 5275–5284.

